# Duplication events downstream of *IRX1* cause North Carolina macular dystrophy at the MCDR3 locus

**DOI:** 10.1101/107573

**Authors:** Valentina Cipriani, Raquel S. Silva, Gavin Arno, Nikolas Pontikos, Ambreen Kalhoro, Sandra Valeina, Inna Inashkina, Mareta Audere, Katrina Rutka, Bernard Puech, Michel Michaelides, Veronica van Heyningen, Baiba Lace, Andrew R Webster, Anthony T Moore

**Author notes:** These authors contributed equally to this work (Valentina Cipriani, Raquel S. Silva). Corresponding authors: Valentina Cipriani, PhD, UCL Institute of Ophthalmology, 11-43 Bath Street, London, EC1V 9EL, United Kingdom; Andrew R. Webster, MD (Res), FRCOphth, UCL Institute of Ophthalmology, 11-43 Bath Street, London, EC1V 9EL, United Kingdom; Anthony T. Moore, MA, FRCOphth, UCSF School of Medicine, Koret Vision Center, 10 Koret Way, San Francisco, California (USA).

## Abstract

Autosomal dominant North Carolina macular dystrophy (NCMD) is believed to represent a failure of macular development. The disorder has been assigned by linkage to two loci, MCDR1 on chromosome 6q16 and MCDR3 on chromosome 5p15-p13. Recently, noncoding variants upstream of *PRDM13* and a large duplication including *IRX1* have been identified. However, the underlying disease-causing mechanism remains uncertain. Through a combination of sequencing studies, we report two novel overlapping duplications at the MCDR3 locus, in a gene desert downstream of *IRX1* and upstream of *ADAMTS16.* One duplication of 43 kb was identified in nine NCMD families (with evidence for a shared ancestral haplotype), and another one of 45 kb was found in a single family. The MCDR3 locus is thus refined to a shared region of 39 kb that contains DNAse hypersensitive sites active at a restricted time window during retinal development. Publicly available data confirmed expression of *IRX1* and *ADAMTS16* in human fetal retina, with *IRX1* preferentially expressed in fetal macula. These findings represent a major advance in our understanding of the molecular genetics of NCMD at the MCDR3 locus and provide insights into the genetic pathways involved in human macular development.

**Abbreviations list:**

aCGH: array comparative genomic hybridization
CNV: Copy number variant
IBD: Identical-by-descent
iPSC: Induced pluripotent stem cell
DHS: DNase hypersensitive site
HH: Homozygosity Haplotype
MCDR: Macular dystrophy region
NCMD: North Carolina macular dystrophy
PCR: Polymerase chain reaction
RCHH: Region with a Conserved Homozygosity Haplotype
SNP: Single-nucleotide polymorphism
SNV: Single nucleotide variant
SV: Structural variant
WGS: Whole-genome sequencing

## Introduction

North Carolina macular dystrophy (NCMD) is a rare autosomal dominant disorder in which there is abnormal development of the macula, a crucial structure of the central retina responsible for central vision and colour perception^1^. Understanding the genetics of rare developmental macular conditions is key for unravelling the mechanism of development of this structure that is found only in higher primates within mammals^1^. NCMD shows fully penetrant inheritance and is considered a non-progressive disorder with a wide range of phenotypic manifestations, usually affecting both eyes symmetrically^2,3^. Phenotypic presentation varies from mild cases with drusen-like deposits covering the macular region but with little or no visual impairment, to severe cases with marked central chorioretinal atrophy and poor vision. Although generally nonprogressive, complications associated with choroidal neovascularization can contribute to visual deterioration.

The molecular genetics of NCMD has been extensively investigated with the disorder being mapped to chromosome 6q16 (MCDR1, MIM:136550) in multiple families of different ethnic origins since the early 1990s^4–10^. A similar phenotype has been assigned to a second locus at 5p15-13 (MCDR3, MIM:608850)^11,3^. Interestingly, several studies reported evidence for ancestral haplotypes at the MCDR1 locus^2,12,13^. Early sequencing studies of the two disease intervals failed to identify exonic disease-causing variants^2,14^. More recently, three novel single nucleotide variants (SNVs) were identified in 11 families at the MCDR1 locus, within a DNase1 hypersensitivity site (DHS), in the noncoding interval between *PRDM13* and the neighbouring overlapping genes *CCNC*/*TSTD3*^15^. Two tandem duplications including the full coding region of *PRDM13,* with some additional upstream and downstream sequence included, were also identified^15,16^. One MCDR3-linked family of Danish origin^3^ was found to carry a 900 kb tandem duplication^15^ that includes the entire coding sequence of *IRX1*. However, duplications of *IRX1* have been observed in several normal individuals from the Database of Genome Variants^15,17^ and the significance of this reported variant is uncertain. Thus, the causative mechanism at the 5p15-13 NCMD locus remains unclear.

In this report we present a combination of genomic investigations in a cohort of 18 NCMD families. The aim of this study was to identify any causative molecular changes and mechanism of disease in these families.

## Results

### Families and brief clinical phenotype description

Eighteen families with phenotypes consistent with a diagnosis of NCMD were included in the study (Table 1 and Supplementary Fig. S1). Four families were previously reported: suggestive linkage at the MCDR3 locus has been recently described for family 1^14^, family 2 was originally reported to be linked to the MCDR3 locus^11^, and families 12 and 13 were linked to MCDR1^7,9^, with family 13 recently found to carry the SNV V2 upstream of *PRDM13*^15^. All families (mostly of small size) showed autosomal dominant inheritance and had at least one individual with Grade 3 disease. DNA samples from a total of 56 affected and 33 unaffected family members were available for genetic analysis.

**Table 1.** Summary of families with two newly reported tandem duplications at the MCDR3 locus and previously identified V2 variant at the MCDR1 locus

| Family number | Family ID | Origin | Phenotype | Experimental procedure | Causative allele change | Nucleotide change | Number of affected family members analysed | Number of unaffected family members analysed | Total number of family members analysed |
| --- | --- | --- | --- | --- | --- | --- | --- | --- | --- |
| 1 | GC19806<sup>14</sup> | Latvian | NCMD | SNP, aCGH, WGS, PCR/Sanger | chr5:4391377-4436535 | 45158 bp duplication | 5 | 1 | 6 |
| 2 | GC15626<sup>11</sup> | British | NCMD | SNP, aCGH, WGS, PCR/Sanger | chr5:4396927-4440442 | 43515 bp duplication | 9 | 8 | 17 |
| 3 | GC15119 | British | NCMD | SNP, aCGH, WGS, PCR/Sanger | chr5:4396927-4440442 | 43515 bp duplication | 4 | 0 | 4 |
| 4 | GC13840 | British | NCMD | SNP, WGS, PCR/Sanger | chr5:4396927-4440442 | 43515 bp duplication | 3 | 0 | 3 |
| 5 | GC19075 | British | NCMD | SNP, WGS, PCR/Sanger | chr5:4396927-4440442 | 43515 bp duplication | 3 | 0 | 3 |
| 6 | GC15475 | British | NCMD | SNP, WGS, PCR/Sanger | chr5:4396927-4440442 | 43515 bp duplication | 1 | 0 | 1 |
| 7 | GC11709 | British | NCMD | SNP, WGS, PCR/Sanger | chr5:4396927-4440442 | 43515 bp duplication | 1 | 0 | 1 |
| 8 | GC16913 | British | NCMD | PCR/Sanger | chr5:4396927-4440442 | 43515 bp duplication | 1 | 0 | 1 |
| 9 | GC4092 | British | NCMD | PCR/Sanger | chr5:4396927-4440442 | 43515 bp duplication | 1 | 0 | 1 |
| 10 | GC23501 | British | NCMD | PCR/Sanger | chr5:4396927-4440442 | 43515 bp duplication | 2 | 1 | 3 |
| 11 | GC15416 | British | NCMD | SNP, PCR/Sanger | chr6:100040987 | G>C (V2) | 2 | 0 | 2 |
| 12 | GC3722<sup>7</sup> | British | NCMD | SNP, PCR/Sanger | chr6:100040987 | G>C (V2) | 12 | 8 | 20 |
| 13 | GC17225<sup>9,15</sup> | French | NCMD | SNP, PCR/Sanger | chr6:100040987 | G>C (V2) | 12 | 15 | 27 |
|  |  |  |  |  |  | **Total** | **56** | **33** | **89** |

Genomic coordinates refer to GRCh37/hg19 assembly. SNP, aCGH, WGS, PCR/Sanger indicate Illumina SNP array, array-based comparative genomic hybridization, whole-genome sequencing and Sanger Sequencing, respectively. Five affected members from five additional NCMD families were also tested for the presence of previously reported SNVs V1-V3<sup>15</sup> and the two tandem duplications found in this study, but none of these affected individuals was found to carry any of the variants.

Figure 1 shows fundus autofluorescence and optical coherence tomography (OCT) images for selected individuals from families 2 and 3. Individual IV:5 from family 2 presents with a well demarcated, relatively symmetrical, bilateral area of macular chorioretinal atrophy, while individual IV:3 from family 3 shows a mild form of disease with relatively symmetrical, bilateral hyperfluorescent drusen-like deposits concentrated in the macular region.

**Figure 1.**
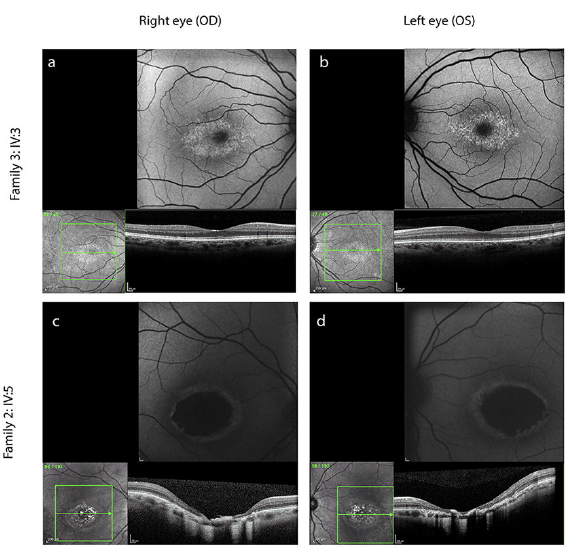
NCMD typical clinical presentation in two selected individuals from family 3 (IV:5) and family 2 (IV:5). Each panel shows fundus autofluorescence and optical coherence tomography (OCT) images. Individual IV:5 (a,b) from family 3 shows a mild form of disease with relatively symmetrical, bilateral hyperfluorescent drusen-like deposits concentrated within the macular region and an otherwise normal OCT. Individual IV:3 (c,d) from family 2 presents with a well demarcated, relatively symmetrical and bilateral area of macular chorioretinal atrophy.

### Haplotype sharing analysis can exclude or suggest genetic mapping at known NCMD loci

Haplotype sharing analysis was carried out using the Homozygosity Haplotype (HH) method^18^ to search for shared identical-by-descent (IBD) chromosomal segments among affected individuals within each family. This analysis was performed in those families for which Illumina single-nucleotide polymorphism (SNP) array data were available for more than one affected family member (families 1-5 and 12-13). The 6q16 MCDR1 locus was excluded in four families, including the two previously MCDR3-linked families 1^14^ and 2^11^ and unreported families 3 and 4 (Supplementary Figs. S2-S5). Family 5 showed evidence for haplotype sharing at many regions across the genome, including both the 6q16 and 5p15-p13 loci (Supplementary Fig. S6). The two previously reported MCDR1-linked families 12^7^ and 13^9^ were confirmed with evidence for a Region with a Conserved HH (RCHH) at the 6q16 locus, and not at the 5p15-p13 locus (Supplementary Figs. S7-S8).

### Two additional NCMD families shown to carry previously reported SNV upstream of PRDM13 at the MCDR1 locus

All families, except families 1-4 for which linkage at the 6q16 locus had been excluded via haplotype sharing analysis, were tested with Sanger Sequencing for the three previously reported SNVs (V1-V3) upstream of *PRDM13*^15^. In addition to the previously reported V2 family 13^15^, two more NCMD families were found to harbour the variant V2 (family 11 and the previously described MCDR1-linked family 12^7^).

### Array-based comparative genomic hybridization (aCGH) uncovers duplications at the MCDR3 locus in three NCMD families

To investigate the MCDR3 locus for the presence of structural variants (SVs), an aCGH experiment using 10,000 probes spanning the region at chr5:11882-10140073 (GRCh37/hg19) was performed in three affected individuals from families 1-3 which did not show linkage at the 6q16 locus (Supplementary Figs. S2-S4). All three families were found to harbour heterozygous duplications of approximately 45kb, downstream of *IRX1* and upstream of *ADAMTS16* (Fig. 2a). The duplications were found to be located in the minimal overlapping regions chr5:4391880-4434888 (GRCh37/hg19) in family 1 and chr5:4397221-4440150 (GRCh37/hg19) in families 2 and 3. These SVs were not seen in 16 control individuals included in the same aCGH experiment, nor were they present in WGS data from 650 individuals with inherited retinal disease^19^ or in publicly available population data (CNV browser)^20^.

**Figure 2.**
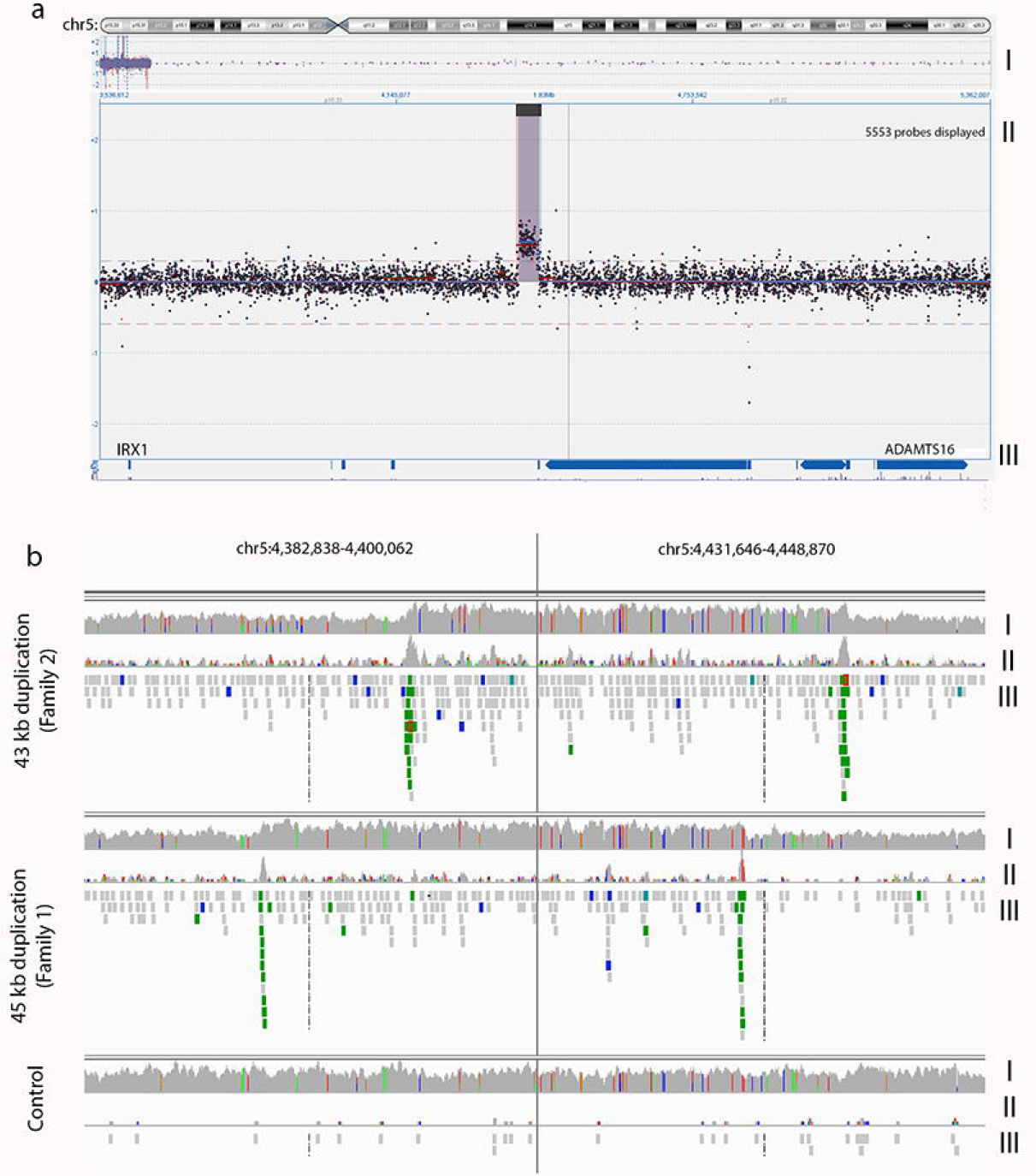
NCMD is caused by intergenic duplication events located between IRX1 and ADAMTS16. (a) aCGH experiment (10,000 probes spanning the MCDR3 locus at GRCh37/hg19 chr5:11882-10140073, panel I) performed in three affected individuals from families 1-3 that were found to harbour heterozygous duplications of approximately 43 kb (panel II) located in a gene desert downstream of *IRX1* and upstream of *ADAMTS16* (panel III), also confirmed by WGS (b) by changes in coverage from concordant and discordant reads (panel I and II, respectively) and identification of chimeric reads, pair-reads with opposing orientation (displayed in green, panel III). Panels are presented with a split view option within IGV. The duplications are located in the overlapping regions GRCh37/hg19 chr5:4391880-4434888 (family 1) and GRCh37/hg19 chr5:4397221-4440150 (families 2 and 3).

### WGS identifies four more NCMD families with duplications at the MCDR3 locus

Thirteen affected individuals from families 1-7 underwent whole genome sequencing (WGS). Graphical visualisation of individual paired-end reads using Integrative Genomics Viewer (IGV)^21,22^ confirmed the presence of heterozygous tandem duplications in families 1-3 (Fig. 2b). Precise breakpoint coordinates were identified from coverage changes, split reads and chimeric reads. Family 1 had a 45158 bp duplicated region (GRCh37/hg19 chr5:4391377-4436535) and families 2 and 3 shared an identical 43515 bp tandem duplication (GRCh37/hg19 chr5:4396927-4440442), overlapping the first identified SV by 85% of the sequence (GRCh37/hg19 chr5:4396925-4436534). Subsequently, members from families 4-7 were also found to carry the same 43 kb duplication.

PCR primers were designed to amplify the novel sequence across the breakpoint between duplicated copies (Table 2, Fig. 3) and used to confirm the predicted breakpoints and assess segregation of the two variants in all available affected and unaffected members of families 1 and 2 (Fig. 3, Table 1 and Supplementary Fig. S1). PCR was then used to genotype the available affected individuals from families 3-7 and confirmed the presence of a band in all affected individuals tested (Table 1 and Supplementary Fig. S1).

**Table 2.**
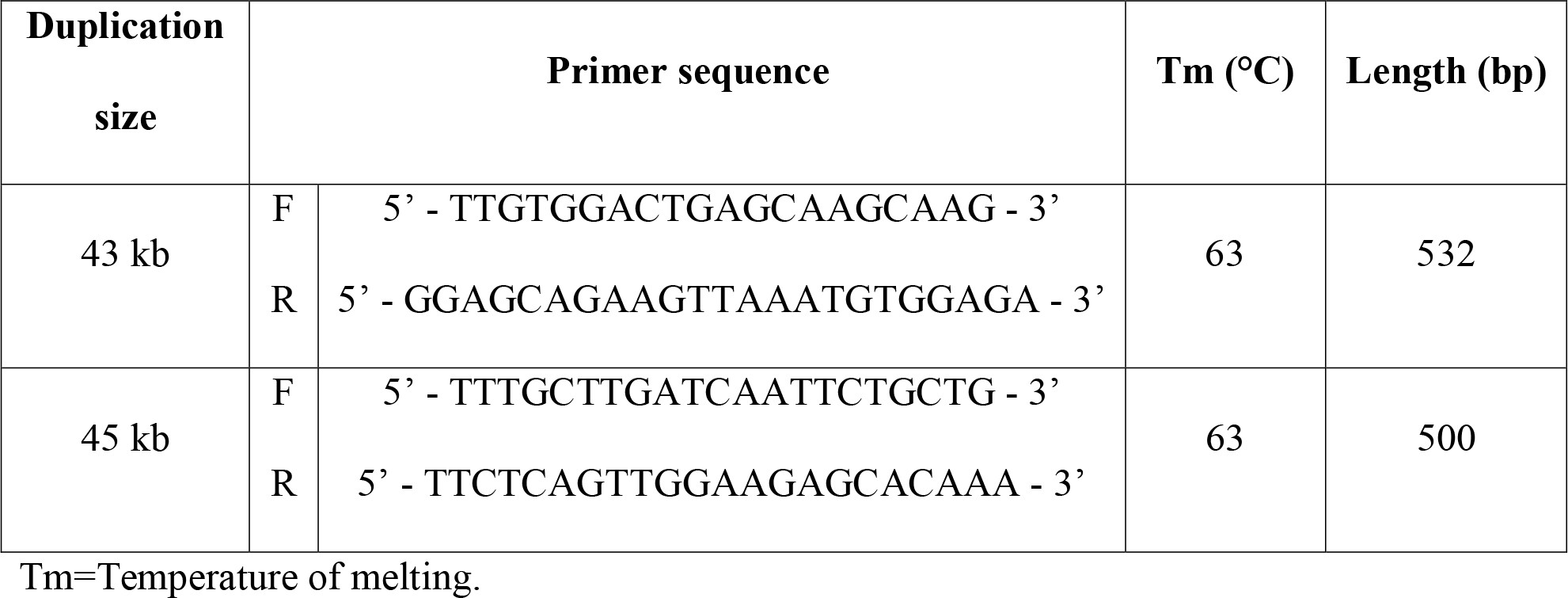
Primer sequences used for the segregation analysis of the two novel MCDR3 duplications identified in the study.

**Table 3.** DHSs active during fetal retina development at the 39 kb shared duplicated region (GRCh37/hg19 chr5:4396925-4436534).

| **Chromosome** | **Start position** | **End position** | **Gestation day (fetal retina)** |
| --- | --- | --- | --- |
| 5 | 4418340 | 4418490 | 74 |
| 5 | 4420820 | 4420970 | 74, 89, 103 |
| 5 | 4418320 | 4418470 | 85 |
| 5 | 4420860 | 4421010 | 85 |
| 5 | 4409260 | 4409410 | 103 |

Gestation day 125 shows no active site at the 39 kb shared region. The fetal retina datasets were available from ENCODE<sup>25</sup>, produced by the Stamatoyannopoulos' laboratory.

**Figure 3.**
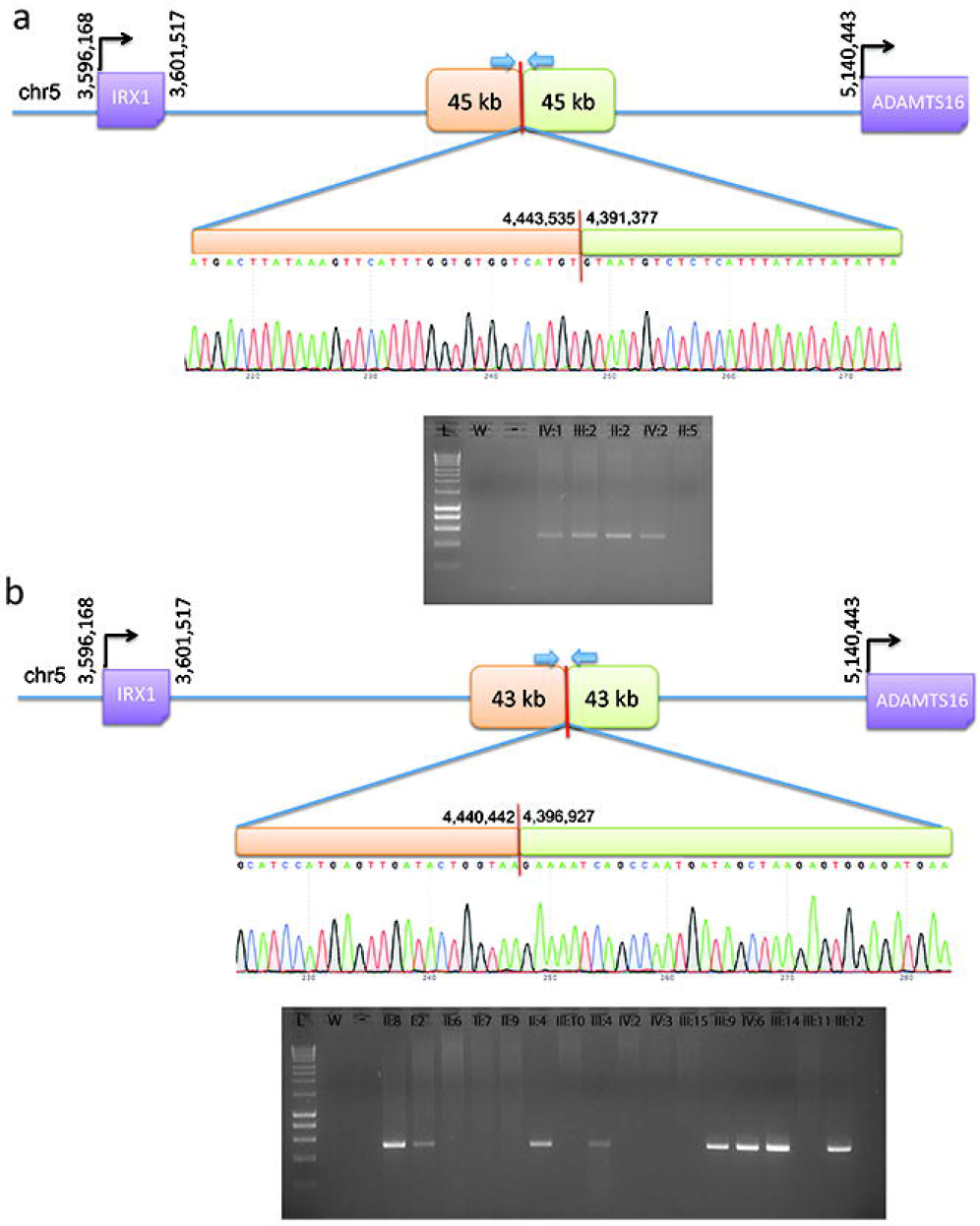
PCR and Sanger sequencing validation of duplication breakpoints and segregation in family 1 (a) and family 2 (b). All available individuals (Supplementary Fig. S1) were tested with primers designed across the predicted breakpoints to generate a unique junction fragment sequence. The exact breakpoint is marked with a red bar; PCR primers are represented with blue arrows. L = ladder; W = water; “-“ = genomic DNA pooled from control individuals.

### Genotyping reveals three additional previously unmapped NCMD families with duplications at the MCDR3 locus

The remaining 8 unmapped families were tested with the established PCR assay for both duplications, and 3 of them (families 8-10) were also found to carry the 43 kb duplication (Table 1 and Supplementary Fig. S1). Thus, a total of 9 not knowingly related families were shown to harbour the same 43 kb tandem duplication at the MCDR3 locus. Five affected members available from the remaining 5 families did not carry either of the two novel duplications.

### Haplotype sharing analysis suggests presence of ancestral haplotypes at the MCDR1 and MCDR3 loci

We hypothesized that finding the same 6q16 SNV and 5p15 duplication with identical breakpoint in 3 and 9 families respectively, could be due to two different shared ancestral haplotypes suggestive of a common founder, in keeping with previous reports on other 6q16 NCMD families^5,12,13,15^. Therefore, haplotype sharing analysis was performed using available Illumina SNP array data from 14 affected individuals in 3 families carrying the 6q16 V2 variant (families 11-13) and 14 affected individuals in 6 families carrying the 5p15 43 kb duplication (families 2-7). Using a cut-off of 2.0 cM and 2.5 cM respectively, the results confirmed that all the genotyped 6q16 individuals collectively shared a RCHH of approximately 2.5 Mb from GRCh37/hg19 coordinate chr6:98962591 (rs150396) to chr6:101468591 (rs1321204) at the MCDR1 locus, and all the genotyped 5p15 individuals collectively shared a RCHH of approximately 0.9 Mb from GRCh37/hg19 coordinate chr5:4327455 (rs155354) to chr5:5210050 (rs1560063) at the MCDR3 locus (Supplementary Tabs. S1-S2 and Supplementary Figs. S9-S10).

## Discussion

We report two distinct heterozygous tandem duplications at the MCDR3 locus in 30 affected individuals from 10 NCMD families. The two novel SVs overlap the previously described duplication found in a single NCMD family of Danish origin^15^ and further refine the 5p15 NCMD locus to a shared region of 39 kb in a gene desert downstream of *IRX1* and upstream of *ADAMTS16* (800 kb and 693.9 kb from the respective transcription start sites, Fig. 4).

**Figure 4.**
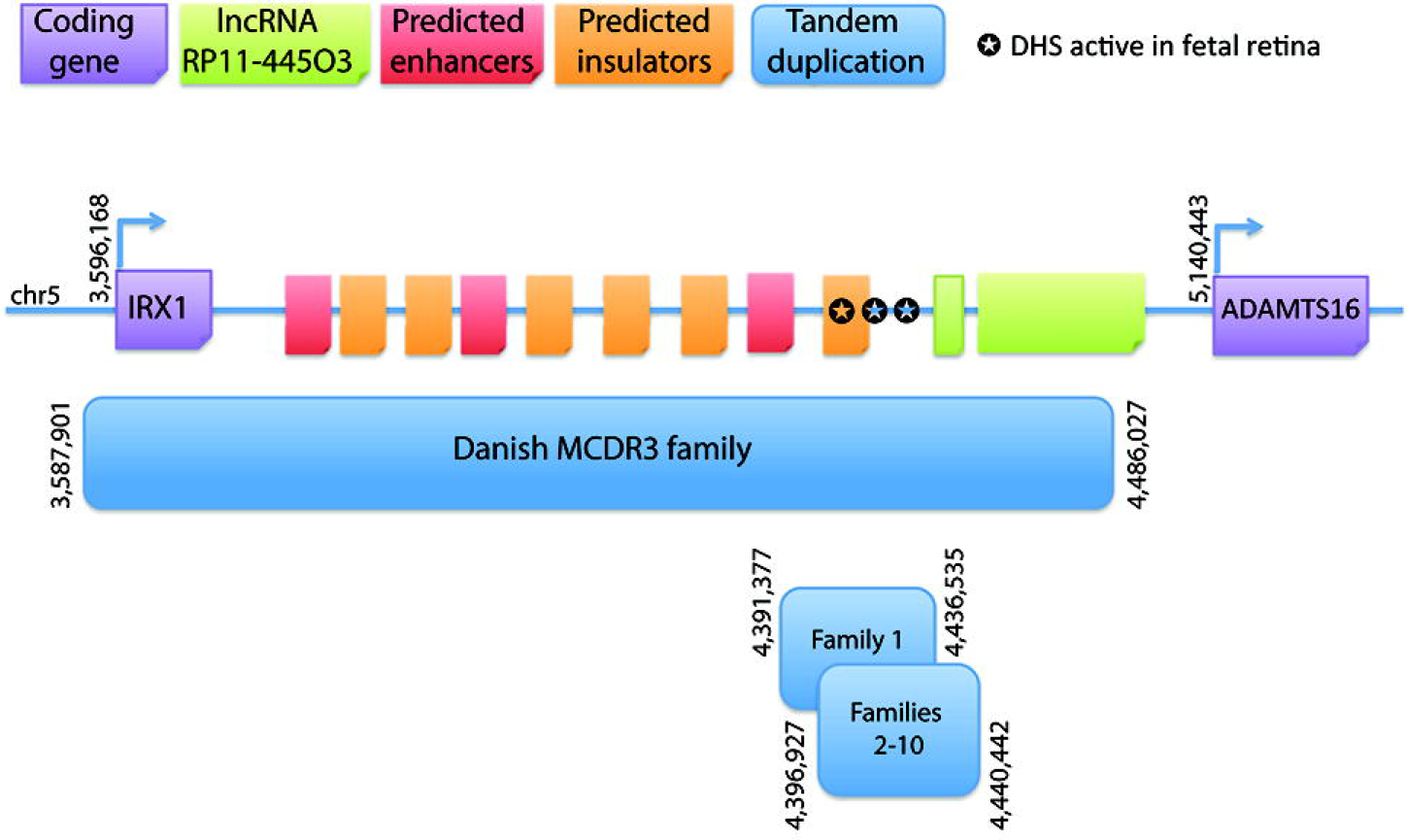
Schematic representation of the MCDR3 locus which is refined to a 39 kb shared genomic region (GRCh37/hg19 chr5:4396925-4436534). The shared sequence between a previously reported duplication and the two novel SVs identified in this study is located in a large gene desert, downstream of *IRX1* and upstream of *ADAMTS16*, 800 kb and 693.9 kb from their respective transcription start sites. Publicly available NGS datasets were queried for informative data on chromatin accessibility and 3 sites were found active from human gestation day 72 to 105 in fetal retina, suggestive of functional acting elements within this site.

We postulated that the 39 kb shared region could harbour *cis*-acting elements that contribute to the fine tuning of gene expression during macular development, affecting target gene expression spatially, temporally and/or quantitatively. Publicly available platforms were queried for informative data on gene expression and chromatin accessibility in relevant tissue types. A dataset screening for gene expression in fetal retina confirmed high expression of *IRX1* at 19-20 weeks of gestation in the macula, and medium expression levels in other regions (Supplementary Fig. S11). In contrast, *ADAMTS16* had medium expression levels throughout the retina^23,24^. Although no role in retinal pathophysiology has been described for *ADAMTS16*, the gene has high sequence similarity to *ADAMTS18* which has been previously associated with retinal disease^32^. Overall, the data suggest that the pattern and/or refined spatial dosage and timing of expression of the transcription factor *IRX1* may be important in macular development. A second dataset provided information on open chromatin conformation using DNase-accessible sequencing in fetal retina tissues at 5 stages from gestation day 72 to 125 (~10 to 18 weeks)^25^. Different sites were identified to be open/active within the 39 kb shared region at four out of five time points (~10-15 weeks of gestation) available during retinal development (Table 3). Interestingly, one of the sites was active during three developmental stages and the remaining four sites were functionally active as two overlapping pairs. At the last time point (day 125, ~18 weeks), all sites were inactive/closed. In the context of human macular development, the sites are active during the period where photoreceptors are proliferating and differentiating^26^; by week 14 of gestation, cells of the central retina exit mitosis^26^, corresponding to the period where DHSs are turning off.

As mentioned, the MCDR1 locus on chromosome 6q16 is associated with variants sited within a DHS, which suggests that aspects of macular development may be highly gene dosage sensitive. Exploring the function and precise target of such regulatory domains in both loci will be essential for understanding the disease mechanism of NCMD and investigating its potential role in the context of normal macular development. The graded expression of *IRX1* and known involvement in retinal development^27,28^, but not *ADAMTS16* in the macular region, suggests that *IRX1* is the probable target of the putative retinal regulatory element which, when duplicated, may cause misregulation of *IRX1*.

Eye development, like other organogenesis processes, requires the precise spatiotemporal and quantitative expression of genes, orchestrated by a complex network of regulatory mechanisms influencing critical transcription factors and other developmental genes. The lack of readily accessible animal or *in vitro* models has hindered detailed understanding of macular development, as this structure only evolved in higher primates among mammals. Recently, disrupted developmental expression of the transcription factor and histone methyltransferase *PRDM13*^29,30^ was suggested as a disease mechanism for NCMD at the 6q16 locus, based on the identification of non-coding SNVs and duplication events residing in an overlapping region upstream of *PRDM13* in many MCDR1 families. Differential regulation of *PRDM13* in eyecups derived from wild-type iPSCs^15,16^ was suggested. However, no causal relationship between the non-coding variants and *PRDM13* expression has been identified.

Despite variable presentation in affected individuals, the NCMD phenotypic spectrum is indistinguishable in patients assigned to either of the two linked loci, MCDR1 and MCDR3. Whether a biological and functional connection between *PRDM13* at the MCDR1 locus and the most likely candidate gene *IRX1* at the MCDR3 locus exists warrants further investigation. iPSC technology and CRISPR manipulation in eye cups from normal and affected individuals may help elucidate the molecular mechanism^33,34^ and the potential molecular links between the two genes. Importantly, the involvement of ancestral variation at both the 6q16 and 5p15 loci (Supplementary Figs. S9-S10) in such a highly penetrant dominant disease is intriguing, with the implication that there may exist a significant number of unrecognized related NCMD families. Full clinical examination reveals a high degree of penetrance, but visually unaffected individuals in whole families may fail to be ascertained.

Finally, the two novel duplications identified in this study significantly further the understanding of the molecular genetics of NCMD at the MCDR3 locus and provide additional effective tools for the molecular diagnosis of NCMD families.

## Materials and Methods

### Families

All families were ascertained at Moorfields Eye Hospital, London, United Kingdom, expect for family 1^14^ (Vision Centre, Children’s Clinical University Hospital, Riga, Latvia) and family 13^9^ (Centre Hospitalier Régional Universitaire de Lille, France).

When possible, retinal imaging was undertaken using colour fundus photography, fundus autofluorescence and OCT imaging. Blood/saliva samples were collected for DNA extraction, genotyping and sequence analyses. The study protocol was approved by the local ethics committees (Central Medical Ethics Commitee of Latvian Republic; NRES Committee London – Camden & Islington) and conformed to the tenets of the Declaration of Helsinki. Written informed consent was obtained from all participants, or their parents, before inclusion in the study.

### Genotyping

Genomic DNA was extracted from whole blood/saliva and genotyped using the Illumina HumanOmniExpress-24 v1.0 beadchip (Illumina, Inc., San Diego, CA, USA). Genotypes were determined using the Genotyping Module in the Illumina GenomeStudio v2011.1 software.

### Haplotype sharing analysis

In order to search for chromosomal segments sharing the same haplotype across affected individuals (within the same family or across different families), the non-parametric HH method^18^ was used for the analysis of those affected individuals that were genotyped with the Illumina array. The HH is a type of haplotype described by the homozygous SNPs only (all heterozygous SNPs are removed) and, therefore, can be uniquely determined on each chromosome. Since affected family members who inherited the same mutation from a common ancestor share a chromosomal segment IBD around the disease gene, they should not have discordant homozygous calls in the IBD region and thus they should share the same HH. The HH approach predicts IBD regions through the identification of RCHHs defined as those regions with a shared HH among affected individuals and a genetic length longer than a certain cut-off value (recommended cut-off for Illumina HumanOmniExpress array is 2.5/3.0 cM for the analysis of one single family).

### aCGH

aCGH was performed at Oxford Gene Technology (OGT) (Begbroke, United Kingdom) using a custom design consisting of 10,000 probes spanning the MCDR3 locus at GRCh37/hg19 chr5:11882-10140073 (approximately 1 probe every 1,000 bp), designed with Agilent e-Array software (Agilent Technologies Inc., Santa Clara, CA, USA), in three individuals from families 1-3 (Supplementary Fig. S1). Sixteen other individuals affected by non-ocular phenotypes were also included in the experiment and used as controls in the analysis. Scanned images of the arrays were processed with OGT CytoSure™ Interpret Software v4.4 using the Accelerate Workflow for calling CNVs. Duplications or deletions were considered when the log_2_ ratio of the Cy3/Cy5 intensities of a region encompassing at least four probes was > 0.3 or <− −0.6, respectively (software default settings).

### WGS and bioinformatics analysis

Whole-genome sequencing was performed using the Illumina HiSeq X10 platform (Illumina, Inc., San Diego, CA, USA), generating minimum coverage of 30X. Reads were aligned to the hg19 human reference sequence (build GRCh37) with novoalign (version 3.02.08). The aligned reads were sorted by base pair position and duplicates were marked using novosort. Discordant reads were marked with samblaster (version 0.1.20) and sent to a separate file for manual inspection of breakpoints using the IGV (version 2.3.61). SVs were manually investigated using the IGV by identifying peaks of discordant reads which were interpreted as breakpoints. The identified duplicated regions were also screened for the presence of common copy number variants using data from the CNV browser^20^ (https://personal.broadinstitute.org/handsake/mcnv_data/) and WGS data from 650 individuals with inherited retinal disease^19^.

### Sanger sequencing validation of duplication events

Segregation analysis of the duplication events identified by WGS was performed using primers (Table 2) designed to span the end of first copy and start of second copy. A graphical representation is shown in Fig. 3. After sequence confirmation with Sanger sequencing, PCR was used to genotype selected individuals from all identified families.

### In silico analysis of duplicated sequences and expression of flanking genes

The Encyclopedia of DNA Elements (ENCODE)^25^ was interrogated for fetal retina datasets of interest. Bed files from DNA-seq datasets (ENCFF249FGP, ENCFF937NUZ, ENCFF401BCF, ENCFF591NRB, ENCFF265ZNN, Stamatoyannopoulos’ laboratory) were downloaded and investigated at the shared duplicated region with R Studio. A second microarray expression dataset on human fetal retina (19-20 gestation week) was queried for the genes of interest^24^ using the platform GENEVESTIGATOR^23^.

## Acknowledgements

The authors would like to acknowledge the ENCODE Consortium and Stamatoyannopoulos’ laboratory for generating the DNA-seq datasets queried in this study, the NIHR BioResource - Rare Disease Consortium (Dr. Keren J Carss and Prof. F Lucy Raymond) for access to CNV data from WGS data, Dr. Gabriela E Jones (University Hospitals of Leicester NHS Trust, Leicester, UK) for help with patient recruitment, Dr. V. Plagnol (UCL Genetics Institute, London, UK) for access to control aCGH data, and UCL Computer Science Cluster and Technical Support (London, UK). This work was supported by grants from the NIHR Biomedical Research Centre at Moorfields Eye Hospital National Health Service Foundation Trust and UCL Institute of Ophthalmology (London, UK), the Research to Prevent Blindness (USA), the British Eye Research Foundation (UK), Fight for Sight (UK), the Macular Society (UK), Moorfields Eye Hospital Special Trustees (UK), Moorfields Eye Charity (UK), the Foundation Fighting Blindness (USA) and Retinitis Pigmentosa Fighting Blindness (UK). RSS is funded through a Fight for Sight PhD studentship granted to ARW and VvH. The views expressed in this publication are those of the authors and not necessarily those of the funding bodies.

## Author Contributions Statement

Study conception and design: ATM, ARW, VvH, VC, RSS, GA.

Patient recruitment and phenotyping: ATM, ARW, AK, MM, BP, BL, SV.

Acquisition of data: VC, RSS, GA, II, MA, KR.

Analysis and/or interpretation of data: VC, RSS, NP, GA.

Drafting of manuscript: VC, RSS.

Critical revision for important intellectual content: ATM, ARW, VvH, GA.

All authors have read and accepted the final version of the manuscript.

## Additional Information

